## Supplementary Materials for "Ultrastructural Visualization of 3D Chromatin Folding Using Serial Block-Face Scanning Electron Microscopy and In Situ Hybridization (3D-EMISH)"

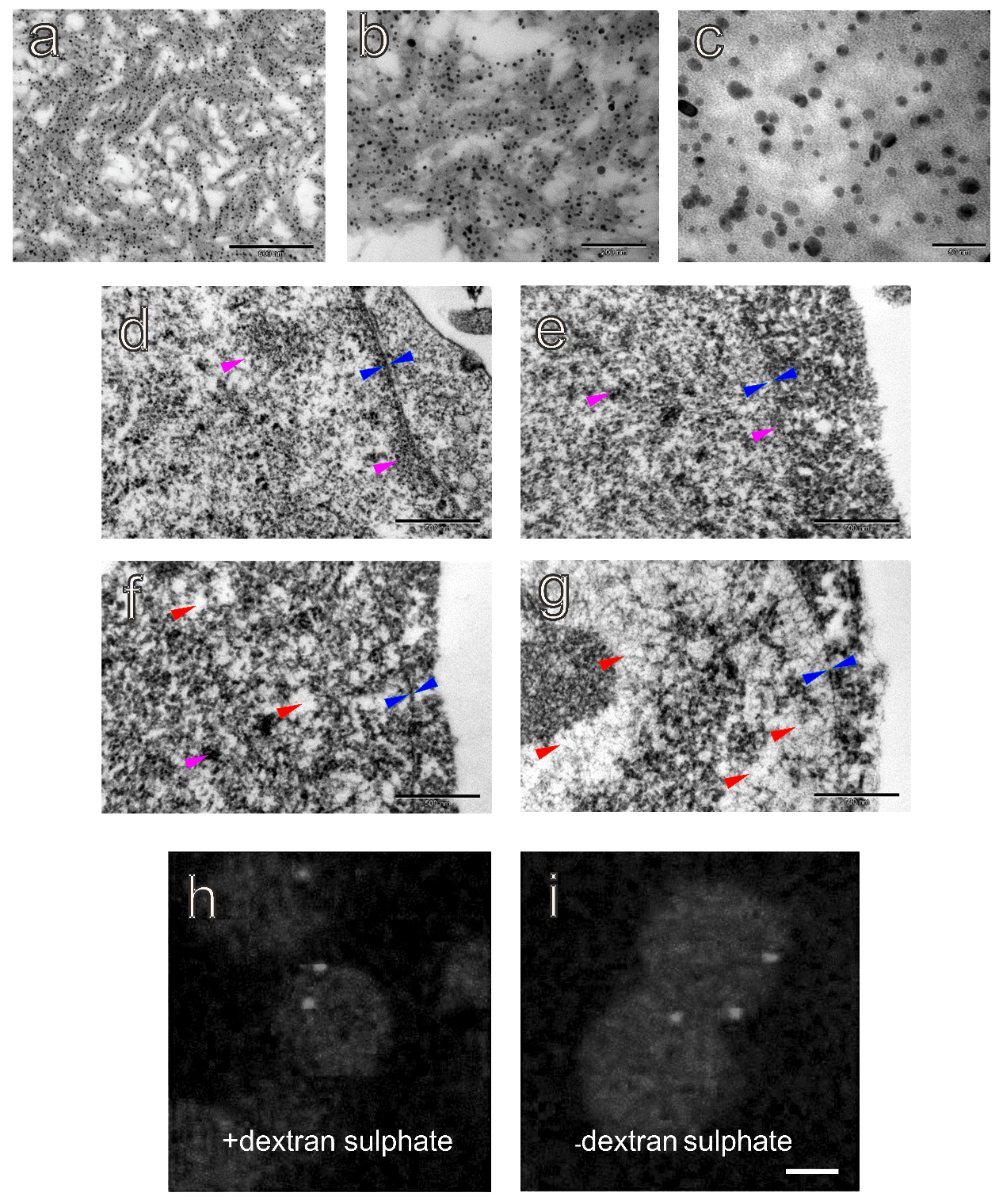

**Supplementary Fig. 1| 3D-EMISH, in situ hybridization effect on chromatin ultrastructure**

We test whether in situ hybridization in 3D-EMISH method alters chromatin ultrastructure, or not. **a-c**, TEM images of streptavidin conjugated silver enhanced fluoronanogold particles used in 3D-EMISH. The particles were added into thrombin and fibrinogen. Three different images (**a**, scale bar, 500 nm, **b**, scale bar, 200 nm, **c**, scale bar, 50 nm, **d**-**g** 500 nm, **h**-**i** 5 µm) show that single silver enhanced fluoronanogold particle conjugated with streptavidin has small enough diameter to improve EM resolution in 3D-EMISH. **d**, TEM image of GM12878 cell nucleus without in situ hybridization procedure as a control case. **e**, TEM image of GM12878 cell nucleus after mock in situ hybridization procedure with the permeabilization, by Triton X-100, without formamide and without dextran sulphate. **f**, TEM image of GM12878 cell nucleus after mock in situ hybridization procedure with permeabilization and formamide/high temperature , but without dextran sulphate. **g**, TEM image of GM12878 cell nucleus after mock in situ hybridization procedure with permeabilization, formamide and dextran sulphate. Blue arrowheads indicate nuclear lamina/envelope and red arrowheads, point out the artifactual changes in nucleoplasmic ultrastructure; magenta arrows indicate condensed chromatin In **e**, compared to **d**, we can see that permeabilization procedure without any other ingredients, causes no serious artifacts in chromatin structure. The addition of formamide/high temperature step to the mock hybridization causes some damage in the form of small, empty spaces in the nucleoplasm and the significant diminution of the condensed chromatin layer beneath the nuclear envelope (compare **e** to **f**). The further addition of dextran sulphate causes the dispersion of chromatin into the network of very thin fibrils and the total loss of the condensed chromatin beneath the nuclear envelope (**g**); scale bars, 500 nm. (**h**, **i**) fluorescence in situ hybridization with (h) or without (i) the addition of dextran sulfate; notice somewhat increased background in (i) compared to (h)

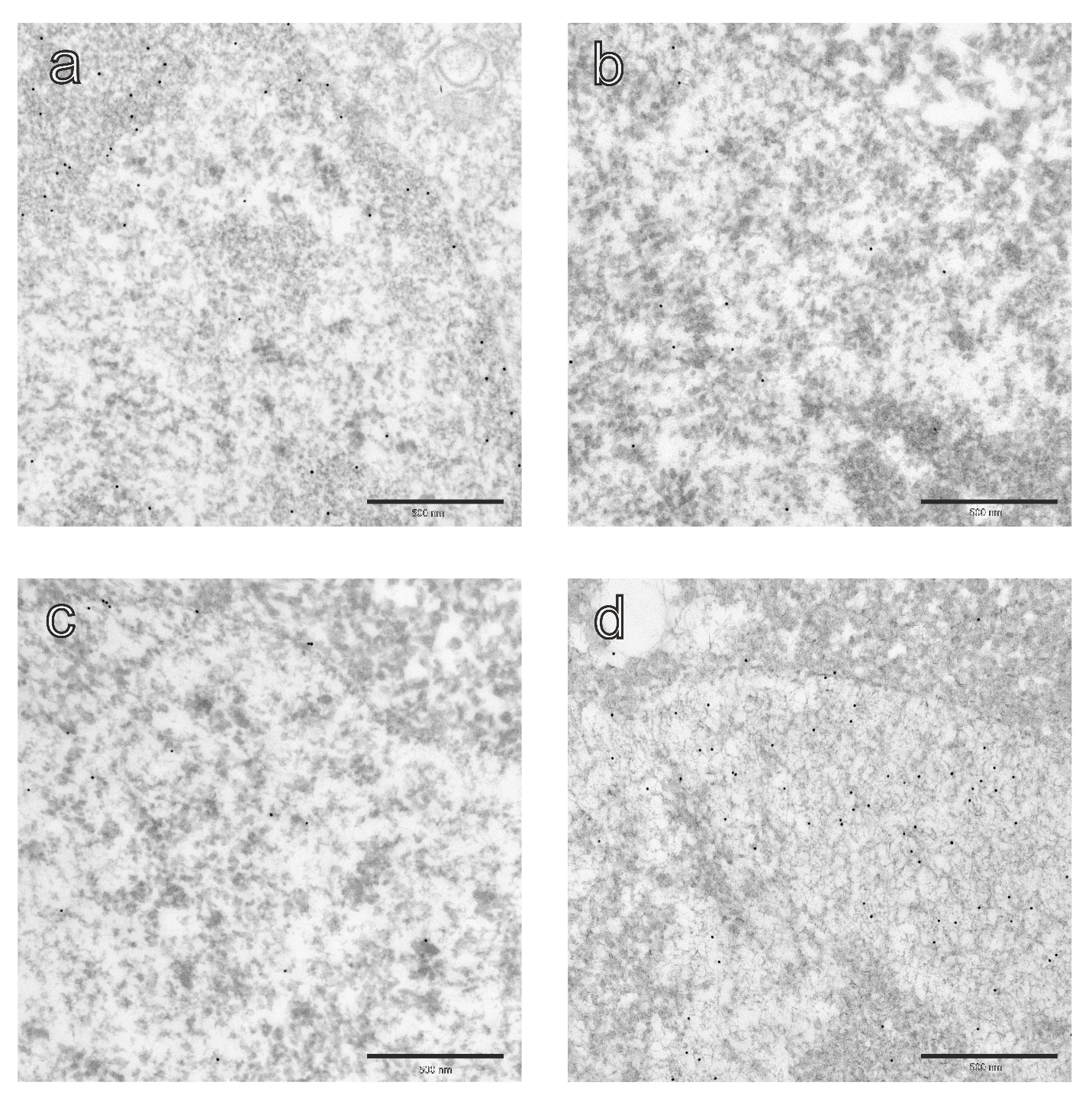

**Supplementary Fig. 2. Specific detection of DNA by hybridization steps with terminal deoxynucleotidyl transferase according to the method by M. Thiry** ().

We determined the impact of factors used in ISH procedure on DNA distribution: **a-d** TEM images of 10nm colloidal gold particles on the sections of en bloc stained GM12878 cell nucleus with uranyl acetate, scale bar 500nm (all panels). **a**, control TEM image of GM12878 cell nucleus without any step of in situ hybridization procedure. **b**, TEM image of GM12878 cell nucleus after mock in situ hybridization procedure comprising permabilization with Triton X-100. **c**, TEM image of GM12878 cell nucleus after mock in situ hybridization procedure with formamide and without dextran sulphate. **d**, TEM image of GM12878 cell nucleus after mock in situ hybridization procedure with formamide and dextran sulphate.

Note the deterioration of chromatin structure, most pronounced after dextran sulfate treatment (**d**). In this case chromatin disperses into a meshwork of thin fibrils containing DNA, indicated by colloidal gold particles, which are found also in the cytoplasm.

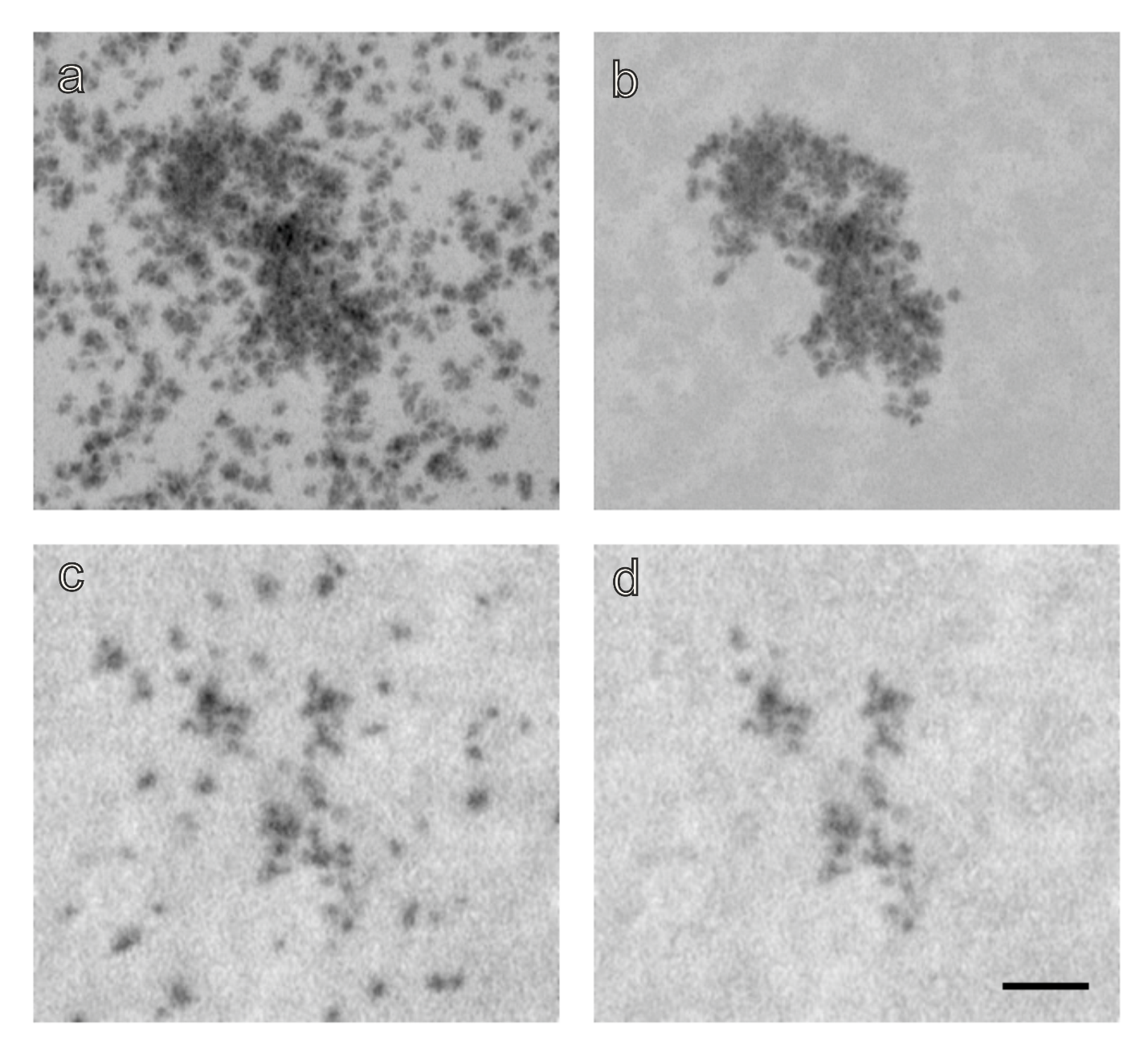

**Supplementary Fig. 3. Filtering of the unspecific background. a,** raw image**,** no filtering; projection of entire stack on a z-plane (minimum signal projection). **b,** image with background filtered; projection of entire stack on a z-plane (minimum signal projection). **c,** raw image**,** no filtering ; single z-section. **d,** image with background filtered, single z-section. Scale bar 320 nm (all panels).

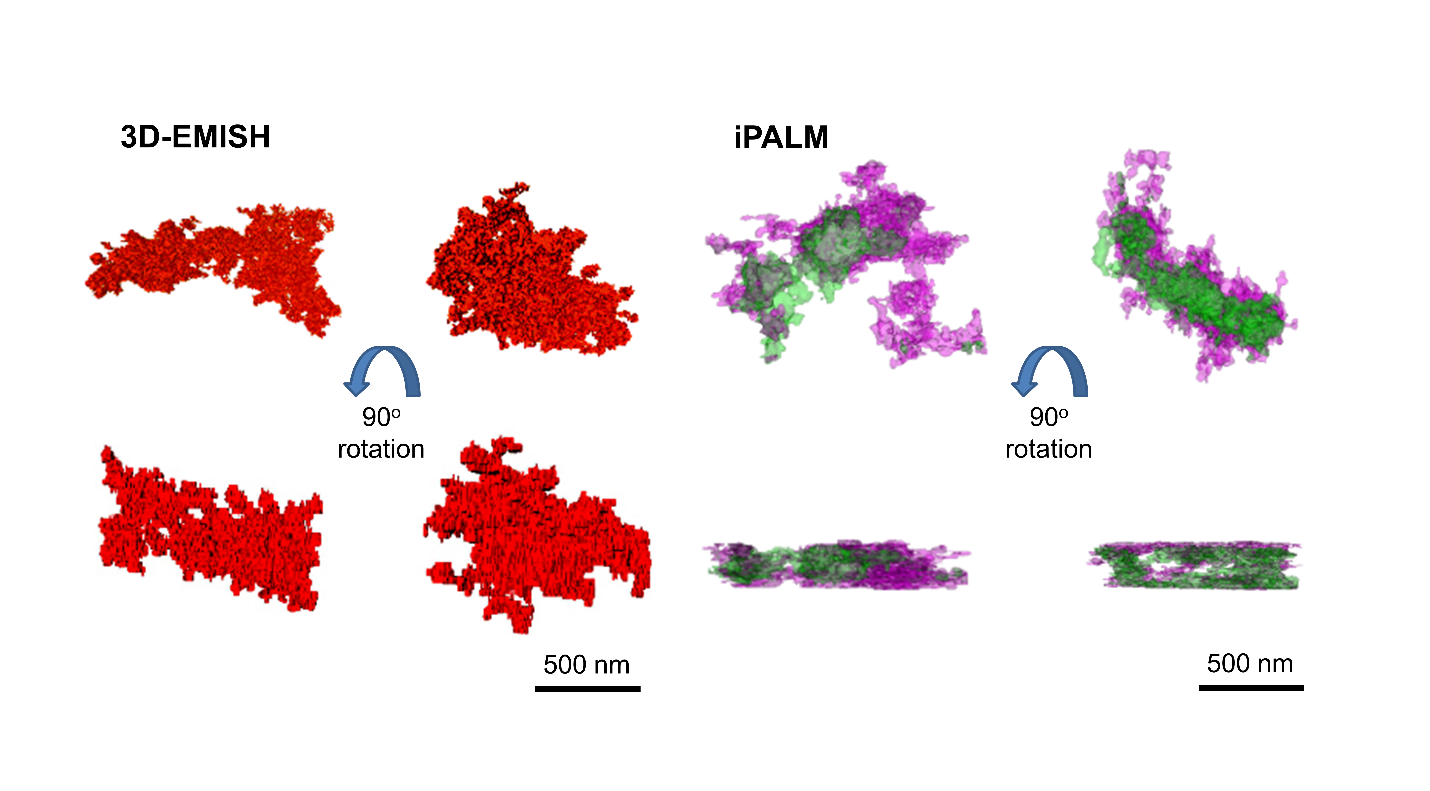

**Supplementary Fig. 3| 3D reconstructed target chromatin structure comparison from 3D-EMISH *vs.* iPALM.**

3D-EMISH DNA probes and iPALM two-color FISH probes were produced by BAC clone, presented in Fig. 2a. 3D reconstructed 3D-EMISH image structures of the targeted genomic region using one color 3D-EMISH DNA probe. 90^o^ rotated structures were presented at bottom to show the whole intact image structures. The voxel size is 5x5x30 nm. 3D reconstructed iPALM image structures of the targeted genomic region using two color FISH. 90^o^ rotated structures were presented at bottom to show the truncated image, because of the iPALM image depth limit. The voxel size is 25x25x15 nm. Scale bar, 500 nm (all panels).

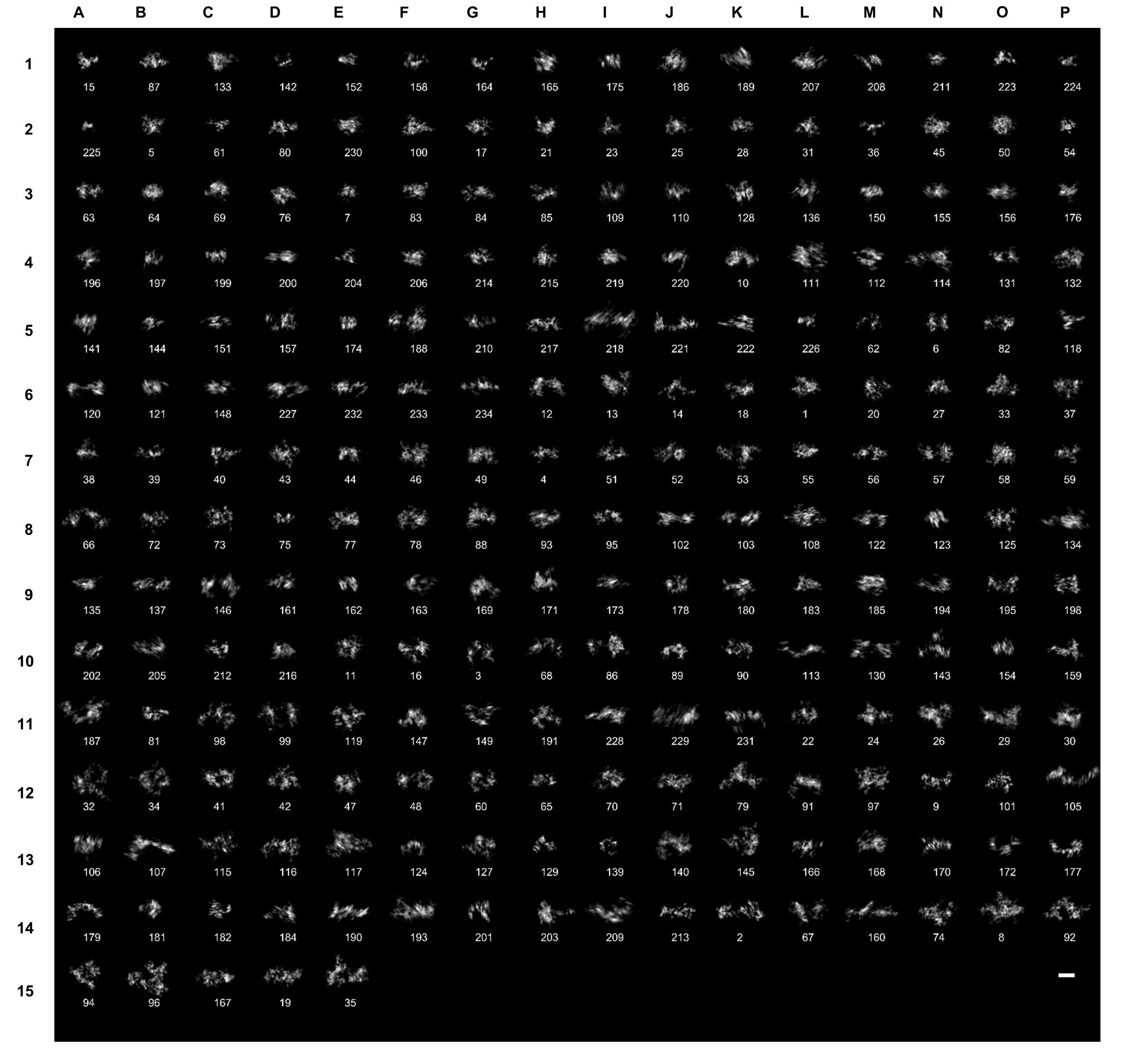

**Supplementary Fig. 4|** Chromatin folding images from 3D-EMISH, maximum projection (229 images collected). Unique structure index number (sID) is assigned to each image. The measurement values of each 3D-EMISH chromatin structure are in the supplementary table; scale bar, 500 nm.

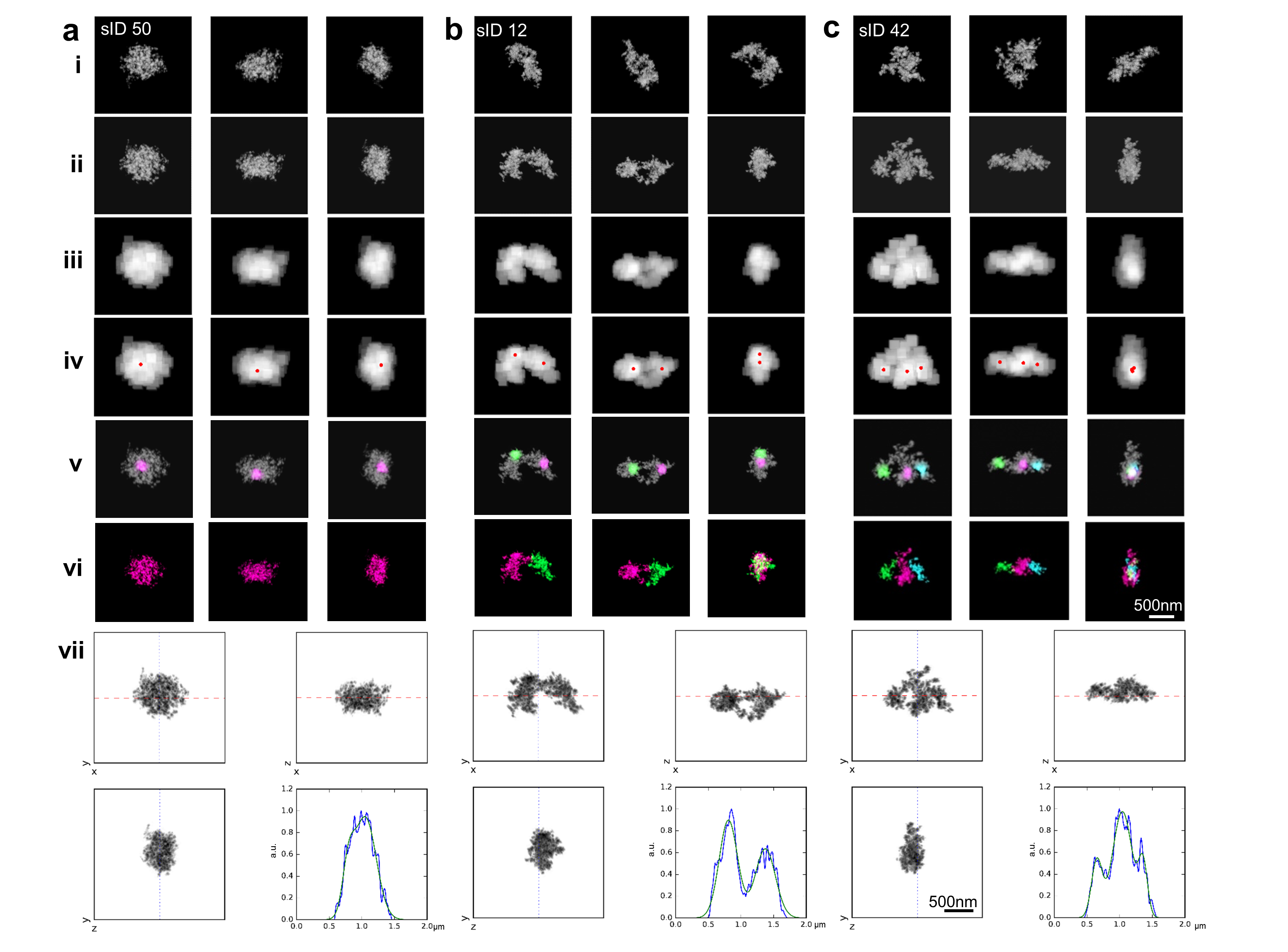

**Supplementary Fig. 5| 3D-EMISH distinctive chromatin folding sub-domain identification procedure**

The identification procedure starts from 3D-EMISH image in the original XYZ coordinate (i), then calculate the principal axes, which orthogonalized the inertia tensor, realigning 3D-EMISH structure in the principal axes coordinate (ii). Not to over-segment domains nor under-segment domains, we smooth out the fine-grain features in 3D-EMISH image by applying smoothing filter to each image voxel, showing the filtered image (iii). Then, we calculate the density center of each domain, representing red dots in (iv). We identify chromatin folding sub-domains in each 3D-EMISH image by diffusing from each local density center point in (iv), until it covers all image structure voxels. (v) intermediate image in diffusion, and (vi) completed image, distinguished each domain by different colors (magenta for the first domain, green for the second domain, and cyan for the third domain). (vii) We validated the chromatin folding sub-domain count in each 3D-EMISH image by projected density signals onto the principal axis, showing one local maximum in sID 50 (**a**), two local maxima in sID 12 (**b**), and three local maxima in sID 42 (**c**), classified as 1 domain, 2 domain, and 3 domain 3D-EMISH chromatin folding structure. We show three different view angles for each image. In (vii), red dashed line is the main principal axis, the blue dotted line is the second principal axis. Then, the EM signal density curve of each image structure is generated by projecting on the main principal axis in xy (top left panel in vii). The gaussian fitted curve (green) is also produced to find local maxima in the density curve.

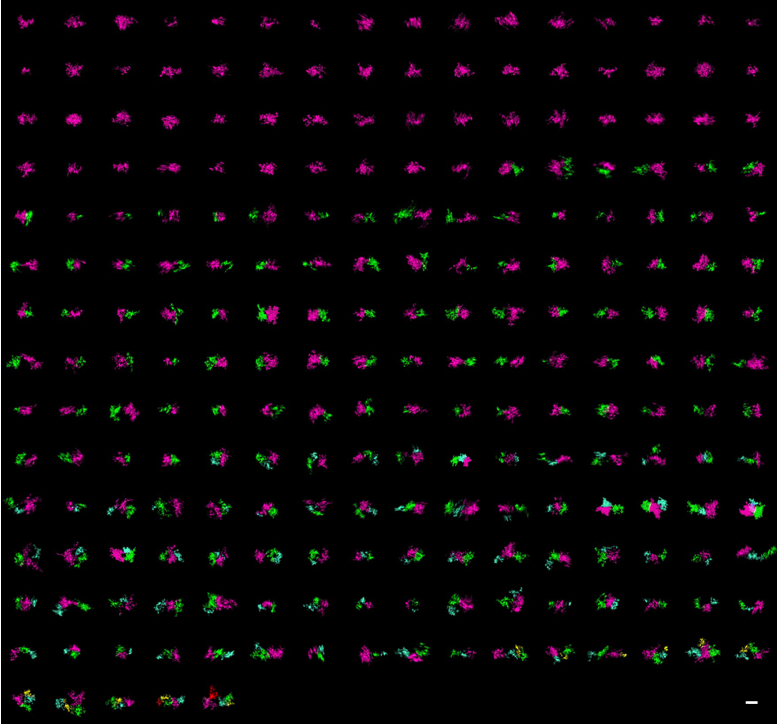

**Supplementary Fig. 6| Distinctive chromatin folding domains identified in 229 chromatin structures.** Identified sub-domains in a structure are highlighted by five different colors: magenta, green, cyan, yellow, and red for the first, second, third, fourth, and fifth domain in structures, respectively. Five domain images are sID 19 (row 15, column D in Supplementary Fig. 3), sID 35 (row 15, column E in Supplementary Fig. 4); scale bar, 500 nm.
